# Exploring SureChEMBL from a drug discovery perspective

**DOI:** 10.1101/2023.11.27.568822

**Authors:** Yojana Gadiya, Simran Shetty, Martin Hofmann-Apitius, Philip Gribbon, Andrea Zaliani

## Abstract

In the pharmaceutical industry, the patent protection of drugs and medicines is accorded importance because of the high costs involved in the development of novel drugs. Over the years, researchers have analyzed patent documents to identify freedom-to-operate spaces for novel drug candidates. To assist this, several well established public patent document data repositories have enabled automated methodologies for extracting information on therapeutic agents. In this study, we delve into one such publicly available patent database, SureChEMBL, which catalogues patent documents related to life sciences. Our exploration begins by identifying patent compounds across public chemical data resources, followed by pinpointing sections in patent documents where the chemical annotations were found. Next, we exhibit the potential of compounds to serve as drug candidates by evaluating their conformity to drug-likeness criteria. Lastly, we examine the drug development stage reported for these compounds to understand their clinical success. In summary, our investigation aims at providing a comprehensive overview of the patent compounds catalogued in SureChEMBL, assessing their relevance to pharmaceutical drug discovery.

## 1. Introduction

Patent documents are legal documents that disclose an invention to the public (https://www.wipo.int/patents/en/). With this disclosure, the holder of a valid patent document generally has the exclusive right to make, use, and sell the invention for approximately 20 years in a given jurisdiction ^1,2^. In drug discovery, researchers explore patent documents to identify competing interests associated with a drug candidate across various organizations, such as pharmaceutical companies, universities, or individuals ^3^. Additionally, patent documents serve as a catalyst for medicinal chemists, empowering them to optimize their drug candidates strategically and ensure their alignment with freedom-to-operate (FTO) zones that may exist outside of the scope of the claimed patent coverage ^4^.

Pharmaceutical-based patenting activity, which mainly covers claims related to therapeutic design, synthesis, and formulation, among others claims, reveals critical information pertaining to the development and prescription of drugs and biologics. In doing so, it serves as a valuable resource for understanding the landscape and dynamics of the pharmaceutical industry ^5–9^. Pharmaceutical patent documents cover two fundamental components: the compound itself and its application ^10,11^. The compound is usually identified in its various forms, such as within a Markush structure, a trade/generic name, etc. Patent documents claim a compound by its structure or even claim a family of structures (based on a scaffold). The chemical structure information is the basis for conducting chemical patent searches by scientists and professionals and is leveraged by commercial vendors in the form of expert software tools and services (eg. CAS-SciFinder) ^12^. The application field(s) of a patent document is usually found in the claims or description sections of the document in text format. A claim’s text description is a legally focused document that often contains specialized terminology and jargon integral to the patent domain. This content plays a crucial role in defining the scope of the underlying patent document, making it a subject of study for patent lawyers and pharma R&D scientists who seek to comprehend the FTO space associated with the patent documents.

Pharmaceutical patent analysis covers a wide range of topics, including patenting trends ^13^, tools for patent protection ^14,15^ and the identification of novel chemical entities for certain treatments or diseases ^16,17^. In a study by Falaguera and Mestres (2021), compounds mined from SureChEMBL, a public patent database, were found to be a collection of starting materials, intermediate products, or pharmacologically relevant compounds (i.e. compounds that target genes or diseases) ^18^. In order to make these compound classifications, the authors employed chemoinformatic methods, such as the matched molecular pair (MMP) analysis ^19^ and the maximum common substructure (MCS) search ^18^, as alternatives to the generic Markush structural searches ^20,21^. Additionally, these methods have allowed for the generation of metrics to assess the chemical novelty and patentability of new compounds ^22^. Despite these efforts, the majority of the aforementioned analyses have been limited to patent documents filed and/or granted in the United States of America (USA). This can be attributed to three main reasons: i) the United States is the world’s largest pharmaceutical market ^23^, ii) the presence of an easy-to-use US-centric public patent database, the United States Patent and Trademark Office (USPTO), which allows for bulk download of patent documents and their metadata ^24^, and iii) the availability of resources, such as the FDA’s Orange Book, which allows for the tracking of drug candidates and their corresponding patent documents through time ^25^. Furthermore, these analyses restrict pharmaceutical patent documents to those tagged with International Patent Classification (IPC) code A61K, an IPC class that includes hygiene-related patent documents in addition to medicinal ones, potentially merging non-pharmaceutical annotations.

SureChEMBL (https://www.surechembl.org/) is an extensive publicly available patent compound data catalogue for the life sciences ^26^. This database identifies compounds, along with other biomedical entities, such as genes and diseases, from patent documents through the use of automated text and image mining pipelines. Furthermore, SureChEMBL keeps an individual record of each extracted compound, associating it with structural information (i.e., SMILES and InChIKeys) and the section of the patent document (i.e. claims, title, description, etc.) where the compound was extracted from. In this study, we aim to investigate the relevance of compounds annotated by SureChEMBL’s pipeline with respect to approved drugs in the pharmaceutical market. Specifically, we applied a medicinal chemistry lens on compounds in SureChEMBL to identify patterns within their molecular scaffolds, as well as the physiochemical properties of patented compounds. Moreover, we assessed the similarity of these compounds to drugs through drug-likeness traits defined by Lipinski (Rule of Five). Rather than limiting the investigation to patent documents found in the United States, as done in all previous methodologies, we broaden our scope to a diverse dataset of patent applications filed and/or granted globally. By doing so, we covered larger IPC patent classes, including information on the medicinal utility of compounds and their formulations. Furthermore, our exploration scrutinizes the availability of compounds described in patent documents and beyond, specifically those annotated by public compound databases. We conclude by delving into the clinical candidate space of the patent compounds in order to understand the success rate of progressing compounds from patent application to clinical practice.

## 2. Results

In the following subsections, we evaluate the chemical space of patent documents found in SureChEMBL, an open-access public patent database. First, we provide a brief statistical summary of the data present in SureChEMBL, with a focus on the country where the patent documents were first registered. Afterwards, we discuss the searchability (i.e. the ability to search for compounds in other databases) of the patent compounds within large chemical databases, namely PubChem, ChEMBL and DrugBank. Next, we review the findability (i.e. the ability to identify the section in patent documents through which the compounds were annotated) of the patent compounds. Following this, we explored the drug-likeness of patent compounds through rules like Ro5 and beyond, evaluated their structural diversity through the Murcko scaffold, and reviewed the presence of structural alerts like Pan-assay interference structures (PAINS) in these compounds. Finally, we briefly discuss the progression of a compound from patent documents to the market through clinical trials.

### Quantitative overview of data in SureChEMBL

From the statistical side, our dataset included a collection of 10 million compounds found in over 1.5 million patent applications (including both granted and non-granted) between 2015 and 2022. Throughout this study, we used the term “patent compounds” to identify those compounds that were captured and annotated by SureChEMBL’s internal pipeline to be associated with a patent application. The patent documents in SureChEMBL are captured across a number of patent offices, namely the USPTO, European Patent Office (EPO), Japan Patent Office (JPO) and World Intellectual Property Organization (WIPO). However, it is worth noting that WIPO usually consists of only filed patent documents and does not have the authority to grant any patent. Additionally, the patent documents in SureChEMBL cover a broad range of IPC classes, such as human necessities (A01, A23, A24, A61, A62B), chemistry and metallurgy-oriented (C05, C06, C07, C08, C09, C10, C11, C12, C13, C14) and physics (G01N), all of which are part of this study.

In SureChEMBL, a patent document is assigned a unique SureChEMBL patent number (SCPN) (for eg. US- 1234567-A1) that consists of a country code (based on the country the patent document was first registered in), followed by a 7-11 digit number, and a patent kind code. In our study, we used the country code in the SCPN to understand the distribution of patent documents across different patent offices. This exploration revealed that patent documents were predominantly filed in the United States and Europe, with 57.3% (24% granted) and 26.6% (11% granted) patents, respectively. Moreover, SureChEMBL also aggregated compounds from Japan (JP), but we found the contribution of these patent documents to be less than 1% ^26^ **(Figure 1A).**

**Figure 1:**
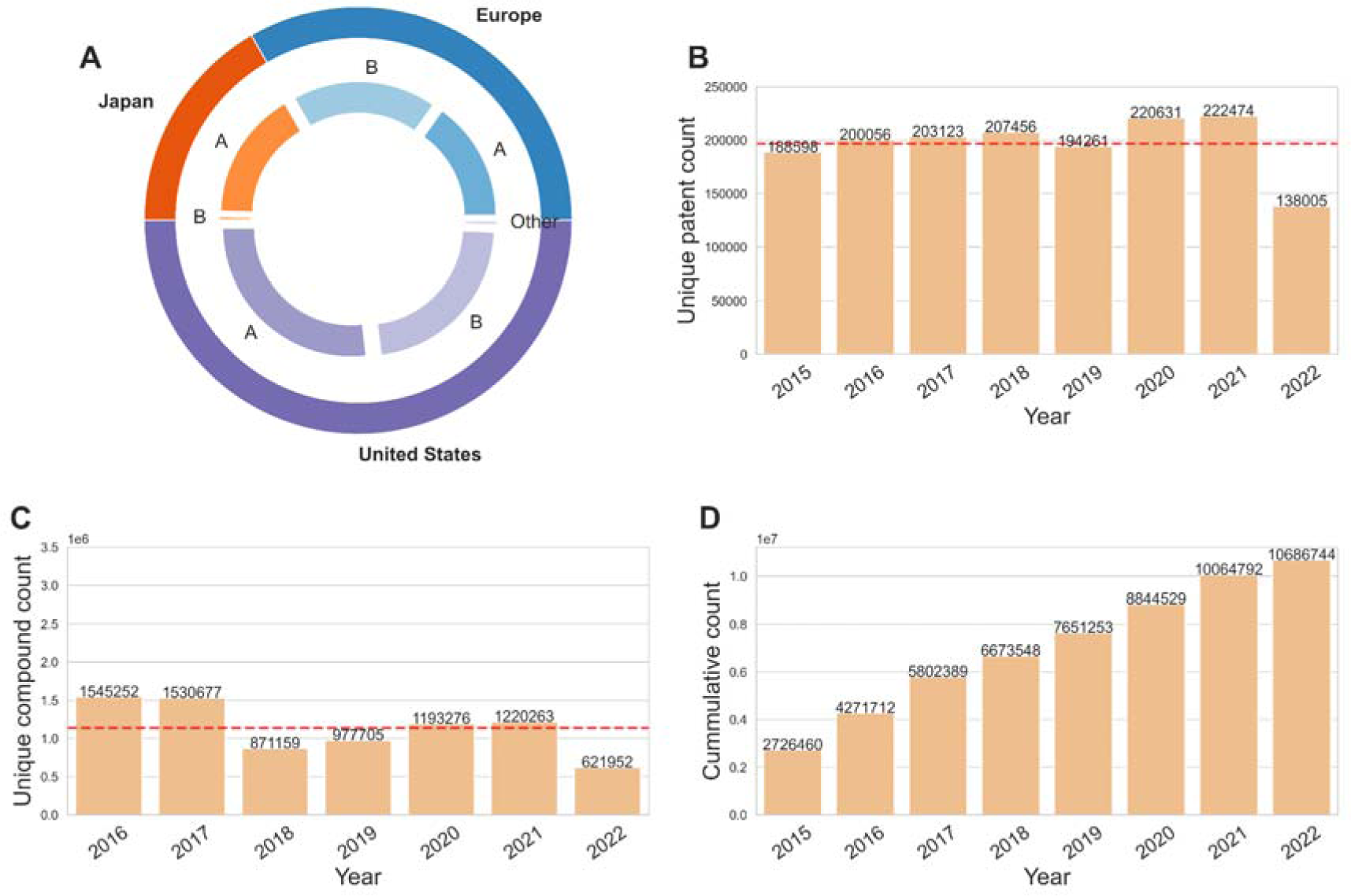
**A) Distribution of patent document types by their patent kind codes across countries. B) Distribution of patent documents filed over the years. C) Distribution of patent compounds over the application years. D) Cumulative compound count over the years**. The red line in subplots B and C indicates the average number of patents and compounds, respectively. Additionally, the counts displayed in subplots B and C are deduplicated counts for patents and patent compounds respectively.

As mentioned previously, each patent document is associated with a “patent kind code” by SureChEMBL. This code is a two-letter alphanumeric code that assists patent officers and reviewers in efficiently distinguishing different kinds of patent documents, such as utility, design, or plant patents. Utility patent documents involve the “discovery and invention of new and useful processes”, design patent documents cover the invention of a “novel design for an article of manufacture”, and plant patent documents cover the scope for “discovering an asexually reproducing variety of plant”. To understand the proportion of these three different patent document types in SureChEMBL, we studied the patent kind codes for each of the registered patent documents in each jurisdiction individually. We identified large proportions of utility patent documents, including both filed (indicated by kind code A*X*) and granted patents (indicated by kind code B*X*) in each of the three countries (i.e. the United States, Europe and Japan), with the prior patent document type being predominant **(Figure 1A)**. Additionally, in the United States, we found the presence of a small proportion of “other” patent document classes, such as design patent documents (indicated by kind code S*X*), reissued patent documents (indicated by kind code E*X*) and plant patent documents (indicated by kind code P*X*) (see **Table ST1** for more details).

Next, we investigated the distribution of patent documents and their compounds yearly. To do so, we collected all the patent documents and the patent compounds in SureChEMBL and distinguished them based on their application number and InChIKeys, respectively. On average, we found that 196,826 patent applications have been filed and patented each year **(Figure 1B and Figure 1C)**. Moreover, an average of 6 compound references were identified per patent document in SureChEMBL. In addition to this, we examined the occurrence of patent compounds in more than one patent document. This analysis illustrated that, of the nearly 1 million patent compounds, 0.2% were associated with multiple patent documents **(Figure SF1).** A detailed investigation revealed that the majority (95%) of these patent compounds were found in less than 5 patent documents **(Table ST2)**.

### PubChem demonstrates highest coverage of patent compounds

A large number of compound-centric biological databases have been established in the past decades ^27–29^. These databases have served various purposes in drug discovery, from identifying the bioactivity of unknown compounds ^30,31^ to the prediction of mechanisms of action ^32,33^, or simply for the virtual screening of drug candidates ^34,35^. While previous research by Joerg Ohms (2022) explored the coverage of patent documents in relation to chemicals in two patent databases (SureChEMBL and Patentscope), the scope of this study was limited to manually comparing a set of chemicals between PubChem Substance and patent compounds ^36^. Thus, to systematically understand the coverage of patent compounds in prominently used chemical databases, we analyzed the structural overlap between the compounds cited in patent documents and those found in three public chemical databases, namely PubChem, ChEMBL and DrugBank. The structural overlap was performed using the InChIKey representation of the compound in SureChEMBL against the three resources.

Upon identifying common compounds across these resources **(Figure 2A)**, two key findings were revealed. Firstly, only 0.02% (2,096 out of 10 million) of the patent compounds were eventually approved for one or more indication areas, according to data extracted from DrugBank, and secondly, PubChem retrieved compounds exhibited the highest overlap (91.5%) with the patent compounds from SureChEMBL. In contrast, ChEMBL demonstrated only a 0.1% overlap with patent compounds in SureChEMBL, an indicator that both resources occupy different chemical spaces. As illustrated in **Figure 2A**, a small percentage (5.5%) of patent compounds were specific to the SureChEMBL database. Among these compounds, more than half were mined from US-based patent documents, while the remaining have been mined from EPO- or WIPO-based patent documents. Additionally, an examination of the annual count of SureChEMBL-specific compounds revealed a gradual decrease over time **(Figure 2B)**.

**Figure 2:**
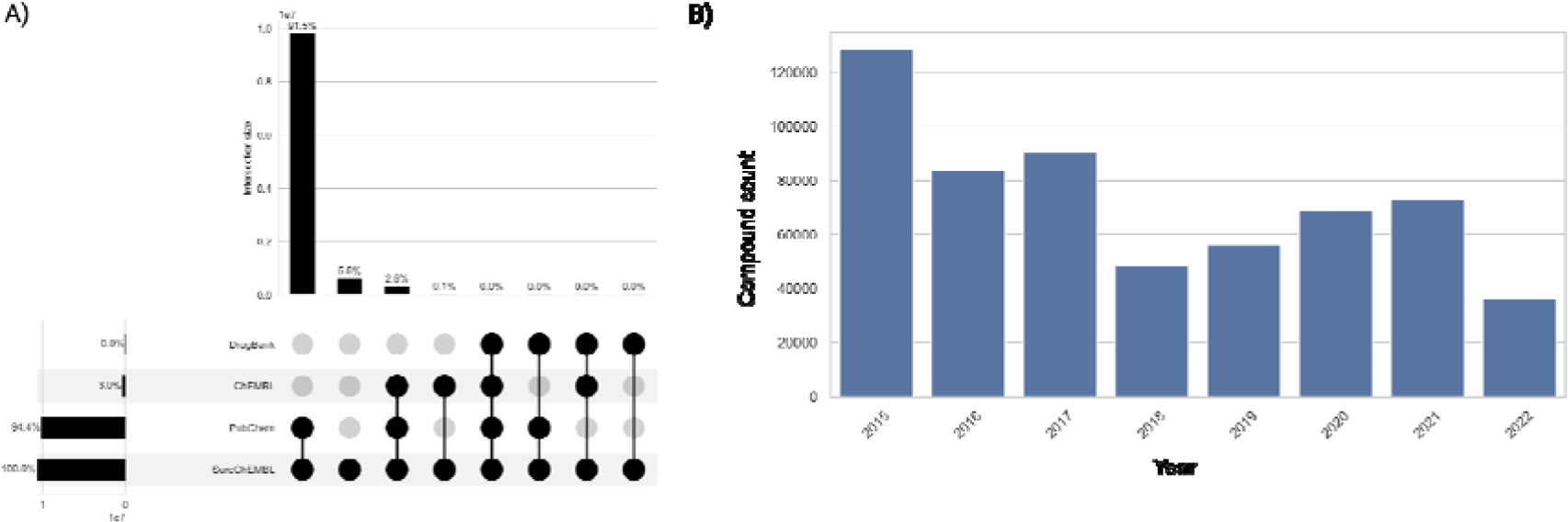
A) Distribution of patent compounds across four chemical resources, namely SureChEMBL, PubChem, ChEMBL and DrugBank. B) Distribution of the proportion of patent compounds specific to SureChEMBL.

### Images as the major source for compound annotation in patent documents

A patent document is a structured document containing sections including the title, abstract, description and claims ^37,38^. Among these patent document sections, determining the location from which a compound was mined can provide insights into the correlation between the compound and the patent’s applicability. For instance, if a compound was mentioned in the description section, it is likely to be associated with prior art (i.e. “referenced” compounds) relevant to the patent document. On the other hand, a compound mentioned in the claims section would likely pertain to the novel invention disclosed in the patent document.

In SureChEMBL, a patent document consists of four sections: title, abstract, description and claims. Along with these specific sections, chemical structure images and molfile (specifically restricted to patent documents collected from USPTO) serve as sources of compound annotation in SureChEMBL. Notably, these later sources (i.e. images and molfiles) were only annotated for patent applications after 2007 ^26^. Together, these six sources of the patent document were utilized for biomedical entity annotations in SureChEMBL using numerous public and proprietary mining tools ^26^. Hence, to provide an overview of the major sources surrounding the annotation of patent compounds, we investigated the sections frequently mined and annotated for compounds in SureChEMBL. We first calculated the average number of sources associated with patent documents in SureChEMBL. This analysis revealed that approximately 31.2% of patent compounds are found in more than one of the six sources. Next, we performed a thorough examination of the sources with regard to the compounds. We found that the description section (with ∼28.08%) of the patent document was the major source of textual data involved in the extraction of patent compounds **(Figure 3)**. As illustrated in the figure, both the additional patent document sources (images with ∼48% and molfiles with ∼15%) were part of the top three sources for data annotations in SureChEMBL.

**Figure 3:**
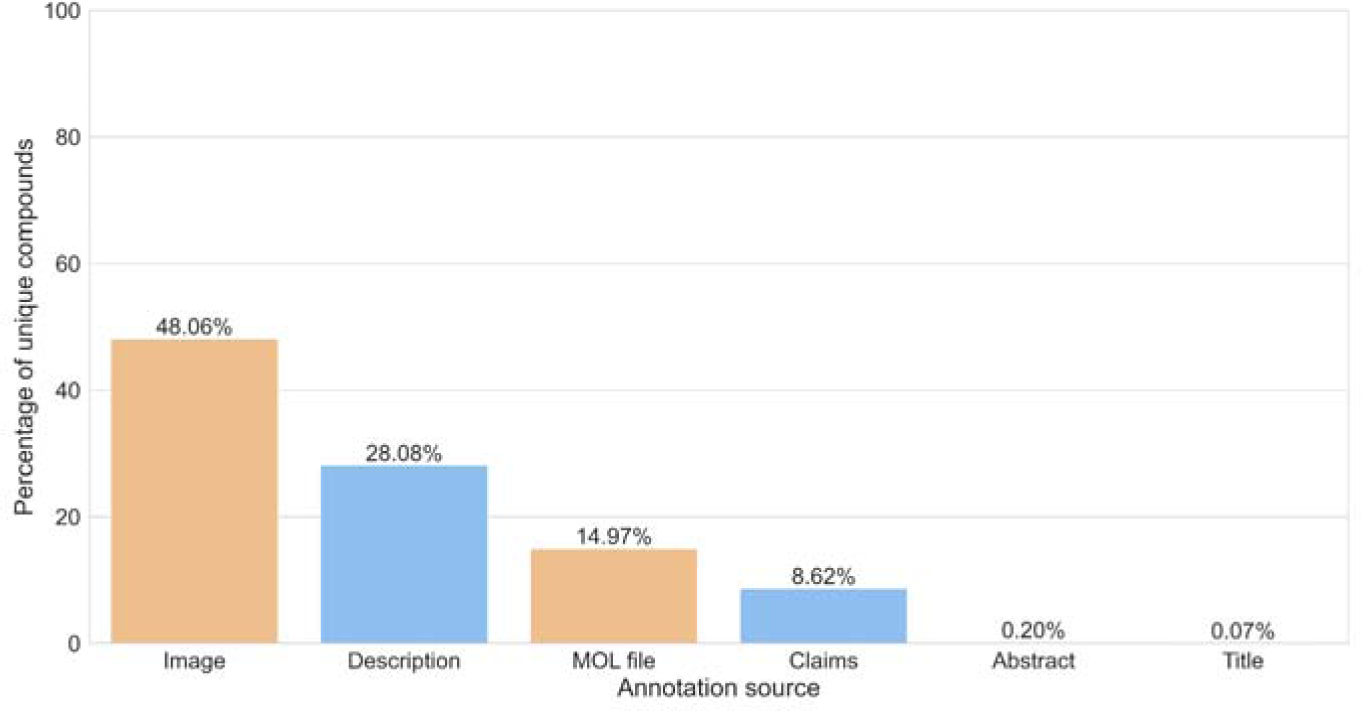
Percentage of compounds annotated from the various patent document sources. Each bar in the figure corresponds to deduplicated compounds annotated specifically from the individual section of the patent document. The textual sections of a patent document (blue) are distinguished from the additional sources for annotation (orange) based on their colour.

### Over half of patent compounds show compliance with Ro5 framework

To improve the efficiency of the drug discovery process, scientists have formulated guidelines or rules based on key determinant properties of compounds of drug likeness. Lipinski (2000) and Veber *et al*. (2002) provided the framework for the Rule of Five (Ro5), depending on physicochemical properties, to enhance the oral bioavailability of a compound ^39–41^. Later, Doak *et al.* (2014), along with other researchers, extended the Ro5 for the oral bioavailability space of drugs, referring to it as the beyond Ro5 (bRo5) space ^42–44^. The criteria of bRo5 supported the selection of cell-permeable clinical candidates that demonstrated good pharmacokinetics (PK) and explored the “undruggable” targets, both of which could not have been possible previously with the Ro5 filtering.

To profile the drug-like space for the patent compounds, we explored the Ro5 and bRo5 space of these compounds. This analysis revealed that in the past decade, 55.46% of compounds complied with the Ro5 framework, and 16.11% compounds did so with the bRo5 **(Figure 4)**. The remaining compounds (28.43%) complied neither with the Ro5 nor bRo5 spaces. Next, we divided the compounds into two categories, approved and non-approved, based on the data in DrugBank. Here, a consistent trend was found, with more than half of the compounds complying with the Ro5 framework, among both approved and non-approved. Finally, we explored the most frequently patent compounds, focusing on the top five based on their prevalence in patent documents **(Figure SF2)**. In the Ro5 class, traditional sugars such as Sorbose (SCHEMBL762) and Mannose (SCHEMBL1812) were found in over 200,000 and 150,000 patent documents, respectively. Essential amino acids like histidine (SCHEMBL3259) and arginine (SCHEMBL1790) were also prevalent, appearing in 150,000 to 200,000 patent documents. On the other hand, in the bRo5 class, we found heparin (SCHEMBL11557), a naturally occurring human metabolite, in more than 95,000 patent documents. Moreover, prominent drugs such as Paclitaxel (SCHEMBL3976), Bleomycin sulphate (SCHEMBL1599) and Doxorubicin (SCHEMBL3243), which are therapeutic drugs for treating cancer and antibacterial drug Streptomycin (SCHEMBL3276) were other patent compounds found in the bRo5 class with occurrence in about 90,000 patent documents. It is important to acknowledge that when dealing with thousands of patent documents associated with a compound, not all of them would be irrelevant. In fact, for a successfully approved active pharmaceutical ingredient (API) with potential, many follow-up patent applications may emerge. These could pertain to its synthesis, specific drug delivery system, novel indication area, or potential combination therapy with another ingredient. Moreover, APIs are frequently cited as prior art in patent documents, underscoring their significance in the pharmaceutical landscape.

**Figure 4:**
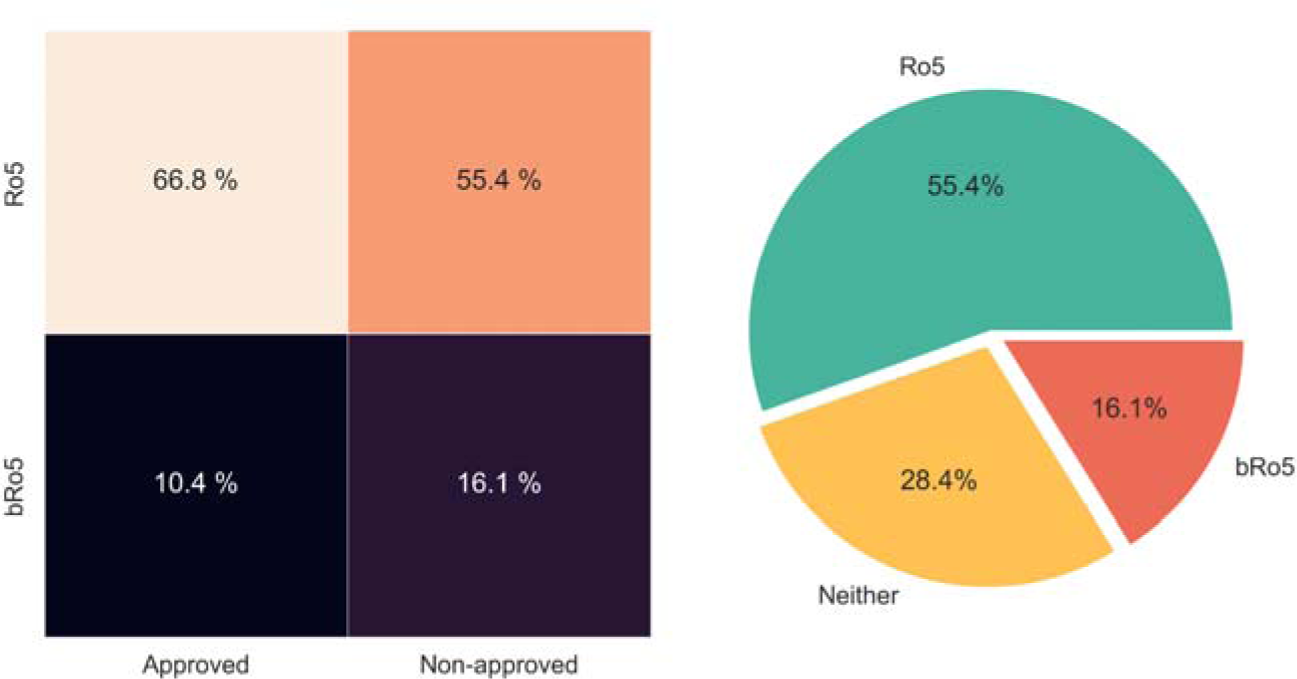
Drug-like compliance of patent compounds. The heatmap on the left shows the percentage of approved and unapproved drugs with regard to the Ro5 frameworks (i.e. Ro5 and bRo5). Here, the compounds that don’t belong to either of the classes have been omitted. On the right is the radial chart demonstrating the percentages of the Ro5 and bRo5 framework compliances of patent compounds. As shown, about 30% of patent compounds do not comply with either of the two frameworks.

### Patent compounds show signs of enhanced chemical structural diversity

Recently, PROteolysis-TArgeting Chimeras (PROTACS) have been identified as novel therapeutics with the potential to progress into clinics ^45,46^. They achieve protein degradation by "hijacking" the cell’s ubiquitin-proteasome system (UPS) and bringing together the target protein and an E3 ligase. Due to their non-adherent Ro5 characteristics, these molecules have not undergone “classical” prior optimization for oral bioavailability ^47^ and CNS penetration ^48^. Additionally, in the past few years, interest has grown in the generation of macrocyclic compounds, those that retain the original scaffold or structure of existing compounds but contain additional functional groups or side chains, allowing for ring-shaped structures ^49, 50^. With the increasing interest in such compounds as potential clinical and drug candidates, we determined the physicochemical properties of patent compounds to check whether the chemical space expansion reflected PROTAC-like and macrocyclic compounds, among others, in recent years.

Table 1 summarizes the average physicochemical characteristics of patent compounds found annually. A gradual increase in characteristic molecular properties (i.e. molecular weight, the hydrogen bond donor and acceptors, LogP, and the number of rings) was observed. These properties, especially molecular weight, were nearing the upper limit of the Ro5 criteria. Specifically, the average molecular weight for patent compounds was between 400 and 500 Daltons, the average LogP was between 4 and 5, the average number of hydrogen bond acceptors and donors were below 6 and 2, respectively, and the average number of rotatable bonds was between 6 and 7. In addition, a consistent increase in the number of rings in patent compounds was also found in the past decade. Our analysis also uncovered some common findings with respect to molecular properties across potential clinical candidates. For example, the number of chiral centers found in patent compounds (summarized in **Table ST3** and **Figure SF3)** mirrors what was recently reported by Scott *et al*. (2022) ^51^.

**Table 1:**
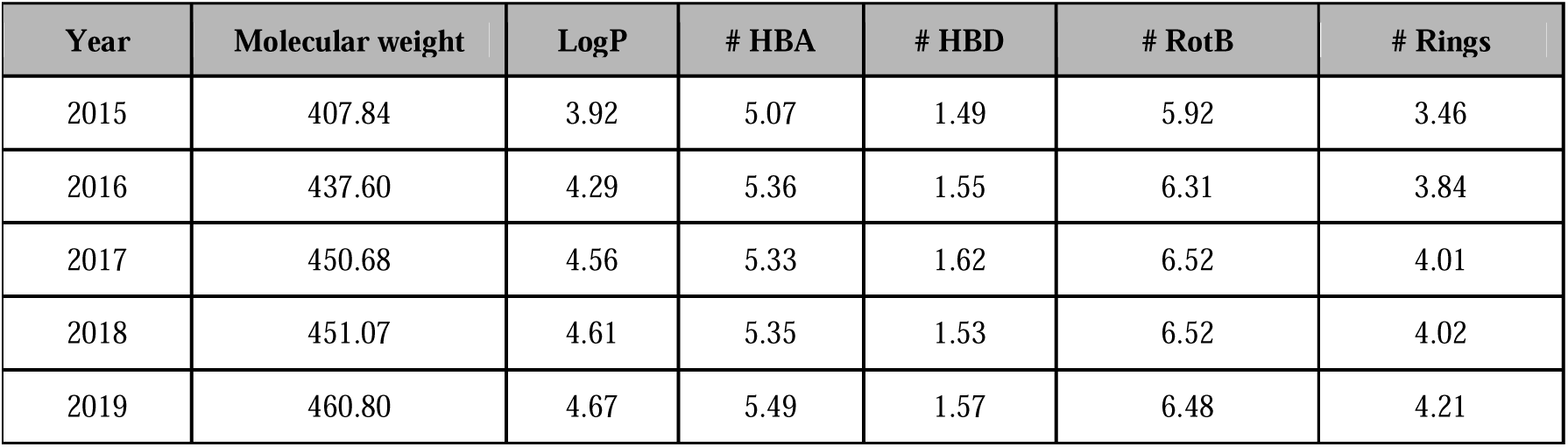
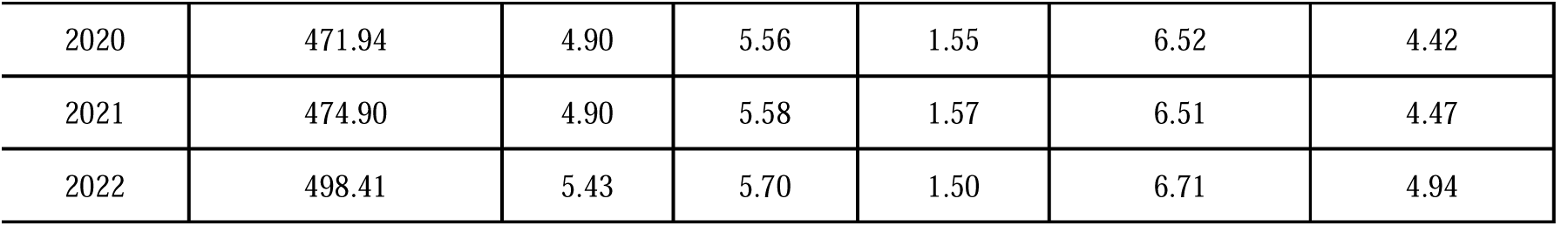
Physicochemical properties of patent compounds. For compounds found in each year, an average of the different molecular properties: i) molecular weights, ii) LogP, iii) the number of hydrogen bond acceptors (# HBA), iv) the number of hydrogen bond donors (# HBD), v) the number of rotatable bonds (# RotB), and vi) the number of any ring (# Rings) were calculated.

Additionally, we were interested in identifying bioactive compounds that could be covalent binders, showed existing polypharmacology, or were reactive in nature. To achieve this, one strategy involved examining the published biological activities and selectivities of the compounds. However, adopting this approach could lead to a very small subset of patent compounds (2,000-5,000), potentially yielding inconclusive results due to the existence of data silos surrounding the publication of biological data in patent documents. Thus, we used an alternative approach to understand the polypharmacological nature of patent compounds. This involved confirming the presence of Pan-Assay INterference Structures (PAINS), which are frequently used in drug discovery to flag and mark compounds that could cause interference in bioassays ^44,52^. Consequently, such compounds that contain one or more PAINS alerts are usually removed during pre-clinical research due to their polypharmacological nature marking them as high risks for off-target effects.

In total, we identified 277 PAINS alerts in the patent compounds. This represents approximately 3.7% of all patent compounds in SureChEMBL that show the presence of at least one of these PAINS alerts **(Figure 5A)**. The most prominent of these PAINS alerts include the presence of azo compounds (Azo_a(324); 18.7%), the presence of compounds involving one aniline and two alkyl groups with either an additional alkyl group (Anil_di_alk_a(478), 14%) or an additional carbon (Anil_di_alk_c(246), 5.7%), the presence of compound containing an indole, a phenyl and an alkyl group (Indol_3yl_alk(461), 8.5%), and the presence of compounds containing catechol substructures (Catechol_a(92), 5.9%). Moreover, aromatic PAINS like quinone also make it to the top of the list with about 5.2% patent compounds **(Table ST4)**. Interestingly, bioassay reagent materials like dyes and Mannich bases also appear in this list of PAINS alerts.

**Figure 5:**
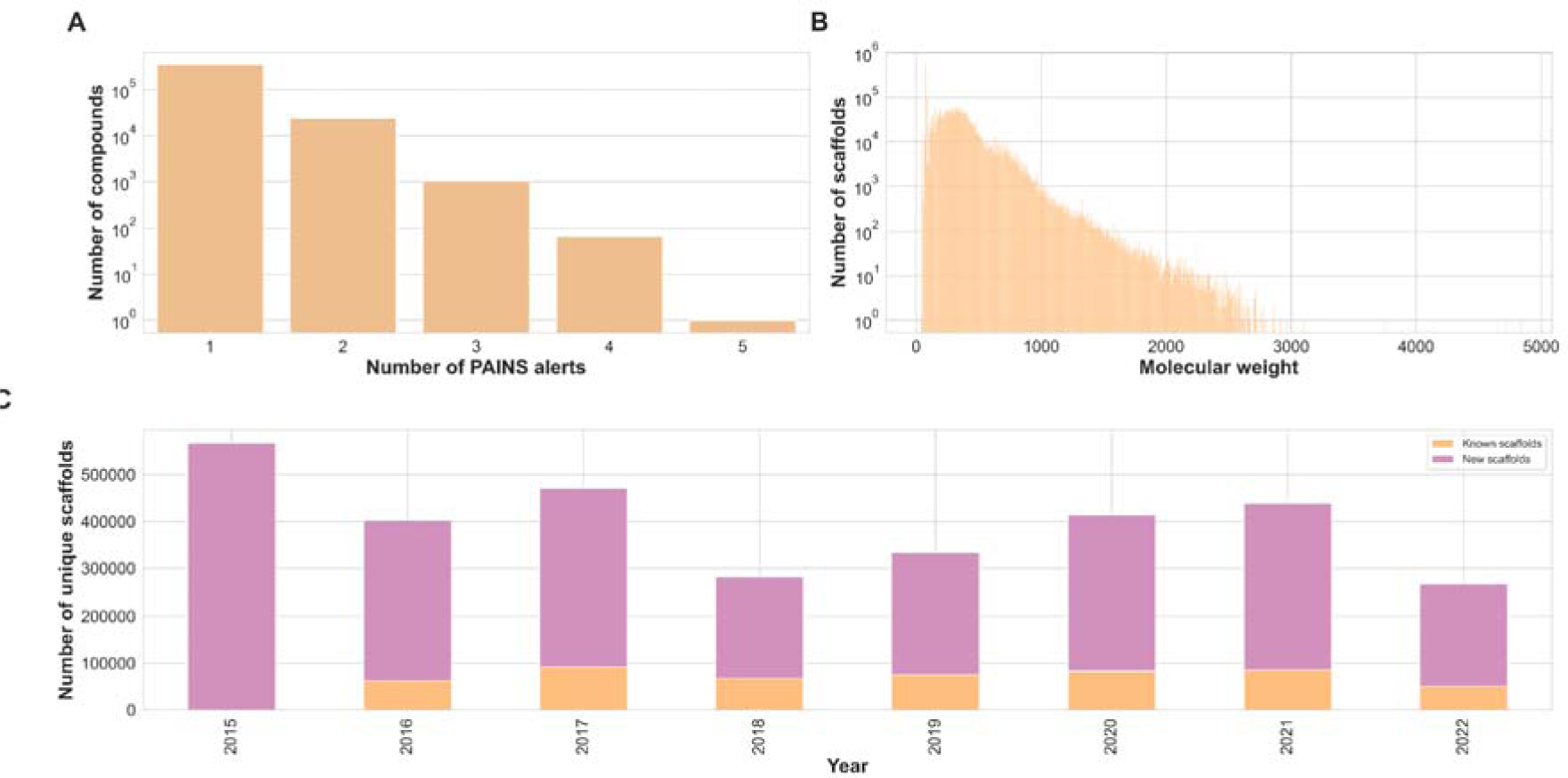
A) Count of PAINS alerts found to be associated with patent compounds. B) Distribution of the molecular weight of the generic Murcko scaffold across the patent compounds. C) Count of unique Murcko scaffolds found per year in patent documents. For each year, we distinguish the “known” scaffolds (orange) from “new” scaffolds (pink) based on the occurrence of the Murcko scaffold SMILES in previous years.

To conclude this exploration, we reduced the compounds to their generic Bemis-Murcko (BM) scaffold to quantify the scaffold diversity of patent compounds. The advantage of using a BM scaffold is two-fold: first, since these compounds are derived from patent documents, mapping them back to their original chemical definition would provide clues on how they were derived, and second, this representation retains the rings and side chains found in the compounds, unlike the graph framework that replaces all heteroatoms to carbon and collapses all bonds to single bonds notations ^53^. This analysis revealed that 3 million distinct scaffolds encompass patent compounds in SureChEMBL. These scaffolds cover a broad range of molecular sizes, spanning from a compact molecule of 38.01 Daltons to a large molecule of 4841.19 Daltons **(Figure 5B)**. The year-wise comparison of the BM scaffold revealed that annually an average of 332,942 new generic scaffolds were patented **(Figure 5C)**. Moreover, trends showcasing a fluctuating number of scaffolds with a recent decline around the COVID-19 pandemic (2021-22) were seen. Tracing back the patent document source (i.e., the section or source from which the compound was annotated) revealed that more than half (55.44%) of the scaffolds were found to be associated with chemical images in patent documents, while only 19.89% were associated with the description section of patent document. Moreover, about 16.62% were found to be extracted from the claims section of the patent document **(Table ST5)**.

In this analysis, it is necessary to acknowledge that reducing the compounds to their generic BM scaffolds would result in the generation of promiscuous compounds like benzene or furan. These common scaffolds might not exactly be found in patent documents but would be part of a larger scaffold shown in these documents. Hence, our approach also unveiled a larger number of such common scaffolds. **Figure 6** summarizes the top ten commonly identified scaffolds in patent compounds. As shown, the majority of these scaffolds have a single ring with heteroatoms causing it to weigh about a few hundred daltons. The most prominent ones include single cyclic scaffolds such as cyclohexane and tetrahydropyran or double-ring compounds like naphthalene and diphenylmethane.

**Figure 6:**
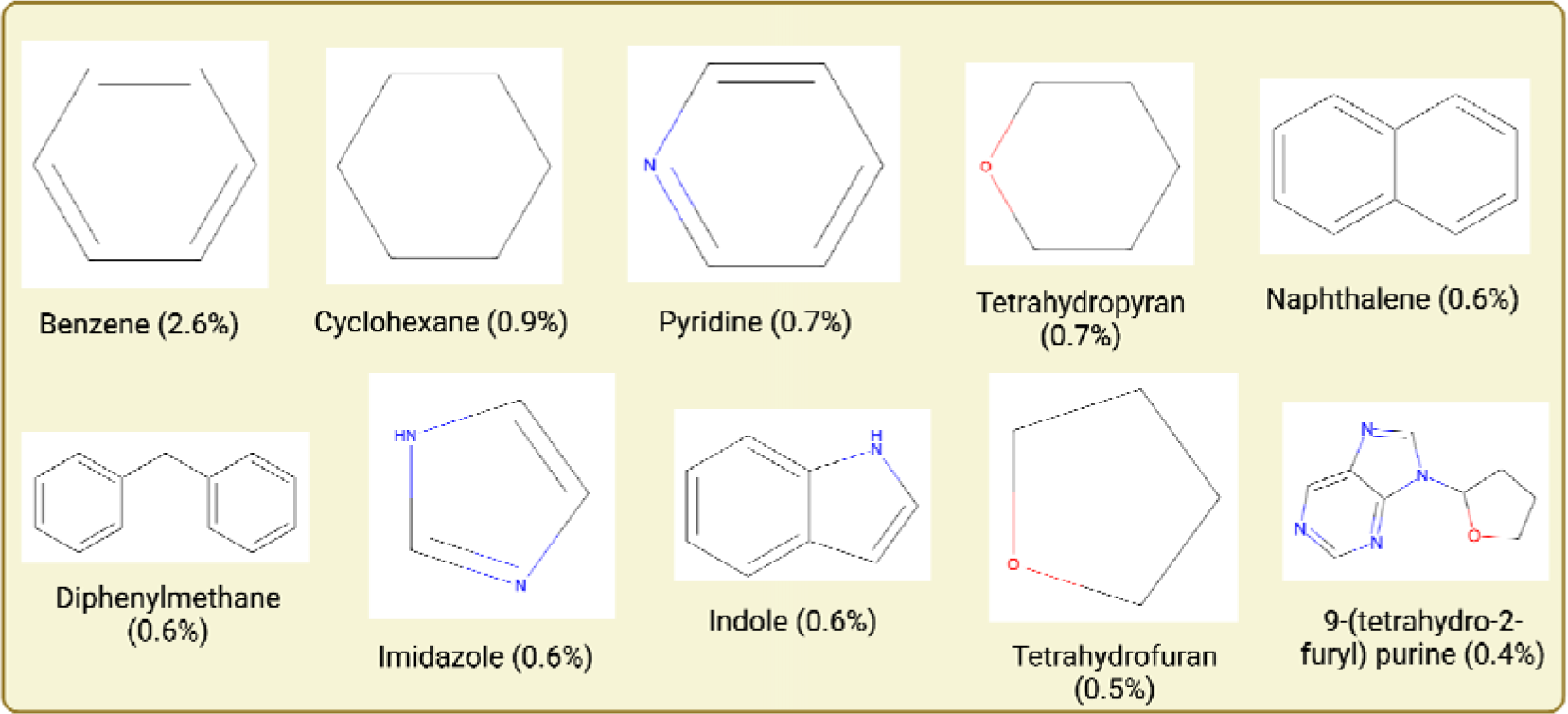
The top Murcko scaffold of compounds and their respective frequency found in patent documents. The percentages only sum up to around 8.2% of the total patent compound scaffolds, a clear sign of the structural diversity within the patent chemical space.

### Tracing a subset of approved drugs back to their patent documents

Pre-established regulations like the Patent Act of 1990 have aided drug proprietors in patenting novel pharmaceuticals or repurposing existing candidates to safeguard inventions under intellectual property laws before their integration into clinical practices ^54, 55^. Consequently, many disparities have arisen between the drug names present in patent documents and their corresponding brand name, posing a challenge in finding associated patent documents ^56^. In the past, successful endeavours were made to link drugs to patent documents by the FDA’s Orange Book and the World Intellectual Property Organization (WIPO) through their Pat- INFORMED tool ^57^. Acknowledging the complexity of the drug nomenclature across the different stages of drug development, we leveraged the chemical representation (InChIKey) of patent compounds to generate an inventory of their corresponding clinical status in humans.

To do so, we started by looking into the intersection of the chemical space of patent compounds with investigational (i.e. drugs that have reached clinical trials) and withdrawn (i.e. drugs that have been discontinued) drugs in DrugBank. We found that only 3,235 of the ten million patent compounds have reached clinical trials, with a mere 0.0008% (85 compounds) falling under the withdrawn drugs category **(Figure 7A)**. In addition, databases such as ChEMBL enable identifying the drug research status (i.e., preclinical, clinical, and approved) of compounds, and hence we leverage this resource to identify the farthest research stage a patent compound traversed to in the drug discovery pipeline. **Figure 7B** depicts that compounds in patent documents are distributed across various research phases, ranging from preclinical to clinical, with the majority concentrated in the preclinical stage. Of these patent compounds, roughly 1% of the drugs were approved for one or more indication areas, according to ChEMBL. Furthermore, 1.6% of the patent compounds had no information (classified as “unknown” by ChEMBL) regarding their clinical stage and were likely to be lost during translation from patent documents to clinics or failed to be captured and annotated by the resource database. Furthermore, Phases 2 and 3 of clinical trials exhibited a greater proportion of patent compounds than Phase 1.

**Figure 7:**
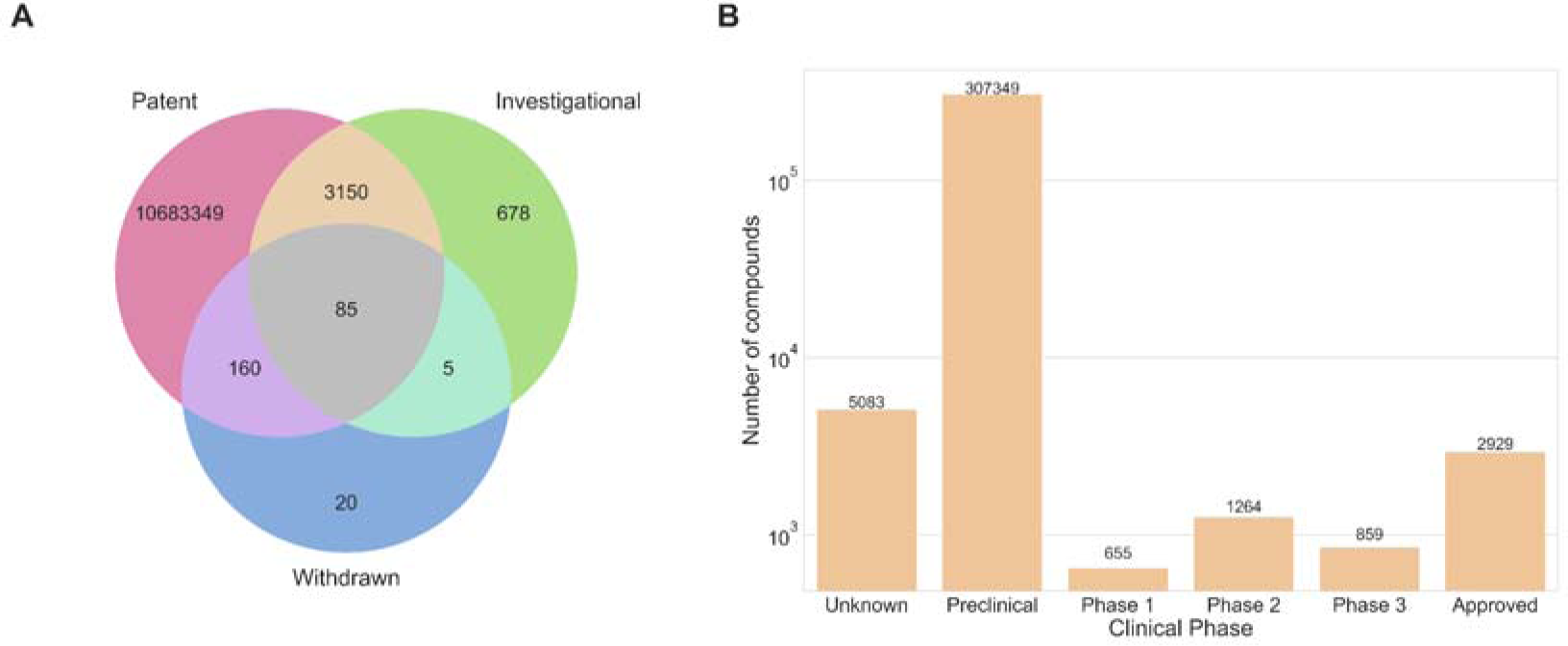
A) Euler figure of the chemical space across the patent compounds, investigational and withdrawn drugs found in DrugBank. B) Distribution of patent compounds in the different clinical phases as per ChEMBL.

## 3. Discussion

Patent documents play an essential role in drug discovery and biological annotation pipelines, such as those that capture a molecule’s image and convert it to SMILES, or those that highlight gene and disease names in the patent documents. This work focuses on patent compounds found in SureChEMBL, a patent database for life sciences, and assesses the annotation quality for drug-like molecules and drug discovery-related documents. To the best of our knowledge, this is the first systematic effort done in this direction with the entire database.

Initially, we started by inspecting the jurisdiction coverage of patent documents in SureChEMBL. As expected, countries such as the United States and Europe had the highest number of patent compounds. In contrast, a very low percentage of patent documents from Japan (through JPO) were present. This low number is confirmed by SureChEMBL, acknowledging their limited access to bibliographic information from Japanese patent documents (i.e. titles and abstracts) provided by the JPO ^26,58^. Furthermore, challenges arise in converting Japanese text to English for the ingestion and storage of biomedical entities in SureChEMBL, thereby exacerbating the limitations. Previously, the absence of machine-translated full texts from which patent compounds could be extracted posed a bottleneck. However, recent advancements in integrated AI annotation within the resource offer potential mitigation for this issue in the future (https://www.ebi.ac.uk/about/news/technology-and-innovation/ai-annotations-increase-patent-data-in-surechembl/). Next, we looked into the quantitative aspect of data in SureChEMBL. A small fraction (0.2%) of compounds were identified in more than one patent document. This could entail the presence of either repurposing patent documents (i.e., a patent application on the reuse of known drugs for a different indication) or compounds annotated from utility patent documents (i.e., a patent application dealing with approved drugs in different pharmaceutical forms or route of administration for specific treatments).

In order to assess the ease with which patent compounds can be identified in the literature, we searched the compound structures across three major compound databases (i.e. PubChem, ChEMBL and DrugBank). In this analysis, we found that a very low percentage of approved drugs (found in DrugBank) were a part of SureChEMBL. This low percentage could be attributed to three reasons: Firstly, patent documents are often crafted to encompass a broader chemical structure-activity landscape than the clinical candidates, thus safeguarding the candidates’ secrecy 46, 59. This is typically achieved through the use of Markush structures in patent claims, allowing for coverage of a wide range of structural variations, including potential drug candidates that may still be unknown at the time of patent filing. Additionally, active pharmaceutical ingredients (APIs) are utilized to extend the applicability of the patent. Secondly, the changes in the nomenclature of the drug candidate as it progresses through the clinical pipeline, obscure its visibility. Despite SureChEMBL’s cross-reference dictionary enabling the retrieval of patent compounds through multiple depictions (e.g., SMILES, IUPAC names, etc.), this limitation arises from inconsistencies in the names used by patent assignees or holders. Additionally, this could be due to the limited information provided by the patent assignees or holders in the patent document (a consequence of the former reason), thus hindering automated systems and pipelines (like SureChEMBL) from accurately recognizing the exact structure of the approved drug. Finally, the third limitation pertains to the use of DrugBank as a proxy for representing the approved drug space. DrugBank, being a commercial database, provides limited information for academic research. Moreover, our results also revealed an analogous chemical space being occupied between patent compounds in SureChEMBL and PubChem. This is unsurprising, given that PubChem leverages the SureChEMBL database to broaden its underlying chemical space. Furthermore, it’s worth noting that in the near future, PubChem could accommodate additional patent compounds through its automated patent annotation pipeline. This pipeline establishes connections between compounds and relevant patent documents referenced in Google Patents. 60. Furthermore, a small proportion of compounds were not found in any public compound resources and were instead confined to SureChEMBL, indicating the presence of compounds from proprietary libraries used by patent assignees for drug discovery.

Following this, we investigated the major sources for compound annotation within SureChEMBL. These sources included known sections of patent documents (i.e. title, abstract, description and claims), images and MOL files. The description section was identified as the prominent textual source for compound annotation, highlighting that the annotated compound could be a part of the primary invention, whether it be related to its synthesis, formulation or application within a specific area. Moreover, it is essential to note that the text within the description section could also be too general, and include compounds such as assay buffers and reactants which are important for the compound stability or assay protocol but not necessarily the main scope of the patent document. Moreover, a small proportion (∼15%) of compounds are also annotated from the MOL file, which is one of the basic files required for compound patent documents filed to the USPTO.

Additionally, we explored the chemical space of patent documents to delineate their structural diversity and drug-likeness space. Naturally, most of the compounds complied with the Ro5 framework for drug-likeness. This was further confirmed by looking at the underlying physicochemical distribution of patent compounds (as shown in **Table 1**). We also identified a small proportion of patent compounds outside the Ro5 space, falling into the bRo5 space; a similar trend was observed in the case of approved drugs in recent years ^42^. Besides this, an increase in the physicochemical properties of the patent compounds was observed, which may have been due to multiple factors, including increasing interest in the development of PROTAC-like and signalling macrocyclic compounds (for e.g., cyclic peptides or cyclic kinase inhibitors) ^61^ or progress with regards to chemical synthesis capabilities ^62,63^. Our exploration regarding the bioactivity of patent compounds led to the recognition of PAINS liable compounds in SureChEMBL. The presence of these assay-interfering compounds is not surprising given that a study by Capuzzi *et al.* (2017) showed that FDA-approved drugs containing PAINS were more active than non-PAINS-containing drugs ^64^. This has indeed led researchers like Senger *et al.* (2016) to question the filtering of promiscuous compounds during the early drug discovery steps ^65^. Nevertheless, the question of whether these PAINS alert patent compounds are problematic (showing false positive results in assays) or innocuous (due to their possible polypharmacology activity) remains unsolved. From our perspective, this finding highlights the notion that the mere identification of compounds from patent documents is not sufficient to identify potential drug candidates. In the case of SureChEMBL, there is a need for performing a medicinal chemistry-oriented filtration to eliminate non-active or activity-interfering patent compounds prior to their downstream utility.

In a similar manner, we also explored the generic BM scaffolds of the patent compounds. This exploration showed a decline in the number of scaffolds generated in the past years. This could be attributed to several reasons, such as exceptional factors like the COVID-19-dependent blockade of some patenting activities in 2022, or structural reasons rooted within medicinal chemistry syntheses pipelines. Lastly, we concluded our analysis by addressing the drug discovery path of a patent drug by looking into its transition from a patent document to post-approval. Here we reported that of all the patent compounds found in ChEMBL, only 1% of the compounds were approved drugs. This is not surprising as an analysis by Brown (2023) showed that hit compounds evolve through the drug discovery pipeline as they undergo structural modification that ensures their entry into clinics ^66^. Moreover, a larger proportion of patent compounds were found in Phase 2 and Phase 3 than in Phase 1. This is attributed to the fact that these trial phases (i.e. Phases 2 and 3) typically yield a larger number of scientific publications, assuming the trials were successful ^67^. In conclusion, our work provides a medicinal chemistry perspective on the chemical landscape formed by patent compounds, thus laying a foundation for the future utility of SureChEMBL. Furthermore, we believe that understanding the state of the art in terms of patent compounds and their scaffolds is crucial for enhancing innovation by exploring novel chemical space while minimizing the risks associated with inadvertently reusing chemical space for specific and undesired target indications.

We acknowledge certain limitations in our analysis that warrant attention. Firstly, our analysis relies on open-source data, which may introduce potential data quality issues. For instance, periodic updates of data sources could lead to temporary gaps, possibly resulting in inaccuracies in our analysis, particularly in areas such as the clinical status of compounds, as discussed in our study. Secondly, we assumed that all patent compounds in SureChEMBL are relevant to drug discovery, as they may pertain to either an indication area or drug formulation. However, this assumption may not always hold true, and it would be preferable to refine our analysis by focusing on a subset of patent IPC codes, as outlined by Gadiya *et al*. (2023) ^68^, to create a more drug discovery-specific patent document dataset. Also, the annotation source in SureChEMBL does not include the context from which the compounds were annotated in the patent document, at least in its data dump. This shortcoming makes it difficult to distinguish compounds that have been “referred” (i.e. prior art) in patent documents from those “claimed” (i.e. novelty). This has eventually led researchers to perform an additional layer of annotation on top of SureChEMBL-extracted patent documents^18,19^. Lastly, we briefly look into drug repurposing patent documents and recognize the potential value in identifying specific indication areas where drug repurposing patent documents are concentrated. This could offer insights into the similarity of MoA across different disease indications for certain compounds.

Discussions on the efficiency of patent documents for drug discovery have been raised numerous times, given their inability to disclose the drug candidate, thereby protecting the compounds’ IP. This study aims to shed light on the utility of compound data in patent documents in the context of drug discovery. By leveraging SureChEMBL as the patent resource and untapping its chemical space, we open the avenue for the use of this resource for chemoinformatic-based models. For example, patent compounds could be used to extend and expand existing chemical datasets by enriching the structure-activity relationship landscape around the lead candidate. Such an approach could be a potential alternative to the generative AI-based approaches, provided that a patent document around the lead molecule exists and has been previously mapped. Hence, SureChEMBL has vast potential that has yet to be mined and leveraged for drug discovery purposes.

## 4. Methods

### Compound databases utilized for compound metadata annotation

To identify compounds annotated within SureChEMBL, we mapped them to three large independent compound data resources, namely PubChem (v.2023) ^60^, ChEMBL (v.32) ^69^, and DrugBank (v.5.1.10) ^70^. The data from these resources was obtained either by querying the REST API service (as in the case of PubChem through PubchemPy), or by downloading a local data dump of the resources (as in the cases of DrugBank, via an academic licence, and ChEMBL, via the SQL database from their FTP server). A compound is said to be the same across two resources provided that an exact match of the InChIKey is present.

Moreover, we leveraged the clinical stage annotation of compounds in ChEMBL (“max phase”) to annotate the clinical phase of a corresponding compound found in SureChEMBL. Additionally, DrugBank was used to validate the compounds that have been approved and distinguish them from those that had been withdrawn. It is worth noting that the commercial nature of DrugBank, while ensuring faster updates than public databases, might limit certain information to premium users.

### Chemical space exploration using physicochemical properties of compounds

The chemical space of compounds was explored using two well-defined and established rules:

a. Ro5 - Extending Lipinski’s Rule of Five (Ro5) ^39,40^, Veber *et al.* (2002) added rotatable bonds (NRotB) and the topological polar surface area (TPSA) as additional features, proposing that compounds should have a TPSA < 140 and NRotB < 12 to enhance the probability of sufficient oral bioavailability ^41^.
b. beyond Ro5/ bRo5 - According to Doak *et al.* (2016), compounds with 500 < MW < 3000 daltons with at least one property beyond the *extended* Ro5 (i.e., LogP > 7.5 or LogP < 0, hydrogen bond donors (HBD) > 5, hydrogen bond acceptors (HBA) > 10, TPSA > 200, and NRotB > 20) fall in this category^42^.

A desalting step using RDKit was performed for the patent compound. In addition to these two rules, we also assessed the chemical space underlying patent documents by conducting a scaffold diversity assessment on compounds derived from patent documents. To accomplish this, we simplified the desalted compounds into their Murcko scaffolds, preserving the generic forms of ring components, linkers and side chains ^43^. These Murcko scaffolds were then used to examine the occurrence of newly registered scaffolds on an annual basis. The generic Murcko scaffold for the patent compounds was generated using RDKit’s “Scaffolds.MurckoScaffold.MurckoScaffoldSmiles()” function. We also identified PAINS alerts within the patent compounds. This was performed using the RDKit library using the codebase from the TeachOpenCADD tutorials ^71^.

## Data Records

### Collecting compound data from patent literature

We used SureChEMBL (https://www.surechembl.org/), an extensive publicly available patent compound data catalogue, as a source for patent documents and metadata ^26^. We obtained all the tab-separated data files (.txt) from the FTP server of the resource (ftp://ftp.ebi.ac.uk/pub/databases/chembl/SureChEMBL/data/map/). The legacy data from 1994-2014 (identified by the file name SureChEMBL_map_20141231.txt.gz) had a different data format, lacking patent information, which complicated the transition from compounds to corresponding patent documents. Consequently, the legacy data was excluded from the analysis presented in the study. Ultimately, the compounds from 2015-2022 were utilized as the patent compound collection for this work.

## Data Availability

The data used in this study can be accessed on Zenodo ^72^. The “Figures” directory consists of all the figures shown in this study. The “Mappings” directory consists of JSON serialized data dumps corresponding to the physiochemical properties, PAINS alerts and Murcko scaffolds for compounds found in SureChEMBL. The “Processed” directory involved the successful mapping of SureChEMBL compounds to external public databases like PubChem and ChEMBL. Finally, the “Raw” directory includes the combined original data dump of SureChEMBL from its FTP server (ftp://ftp.ebi.ac.uk/pub/databases/chembl/SureChEMBL/data/map/).

## Code Availability

The Python scripts and Jupyter notebooks supporting the conclusions of this study can be accessed and downloaded via GitHub (https://github.com/Fraunhofer-ITMP/patent-clinical-candidate-characteristics). The repository is structured into “Data” and “Notebook” sections. The “Data” section is the exact replica of the Zenodo dump mentioned previously. The “Notebook” section includes all the analyses presented in this study organized based on the sections within the results.

## Authors’ contributions

YG conceived the work. YG and AZ contributed to the ideation. YG and SS performed the analysis. MHA, PG, YG and AZ have written the manuscript. All the authors have reviewed and approved the final manuscript.

## Acknowledgements

We would like to thank the authors of the public resources used in our work for making their datasets available to the scientific community. The REMEDi4ALL project has received funding from the European Union’s Horizon Europe Research & Innovation programme under grant agreement No 101057442. This work reflects only the authors’ view, and the EC is not responsible for any use that may be made of the information it contains. We would also like to thank Dr. Sarah Mubeen for helping us improve the language and content of the manuscript.

---

**Supplementary Figure SF1:**
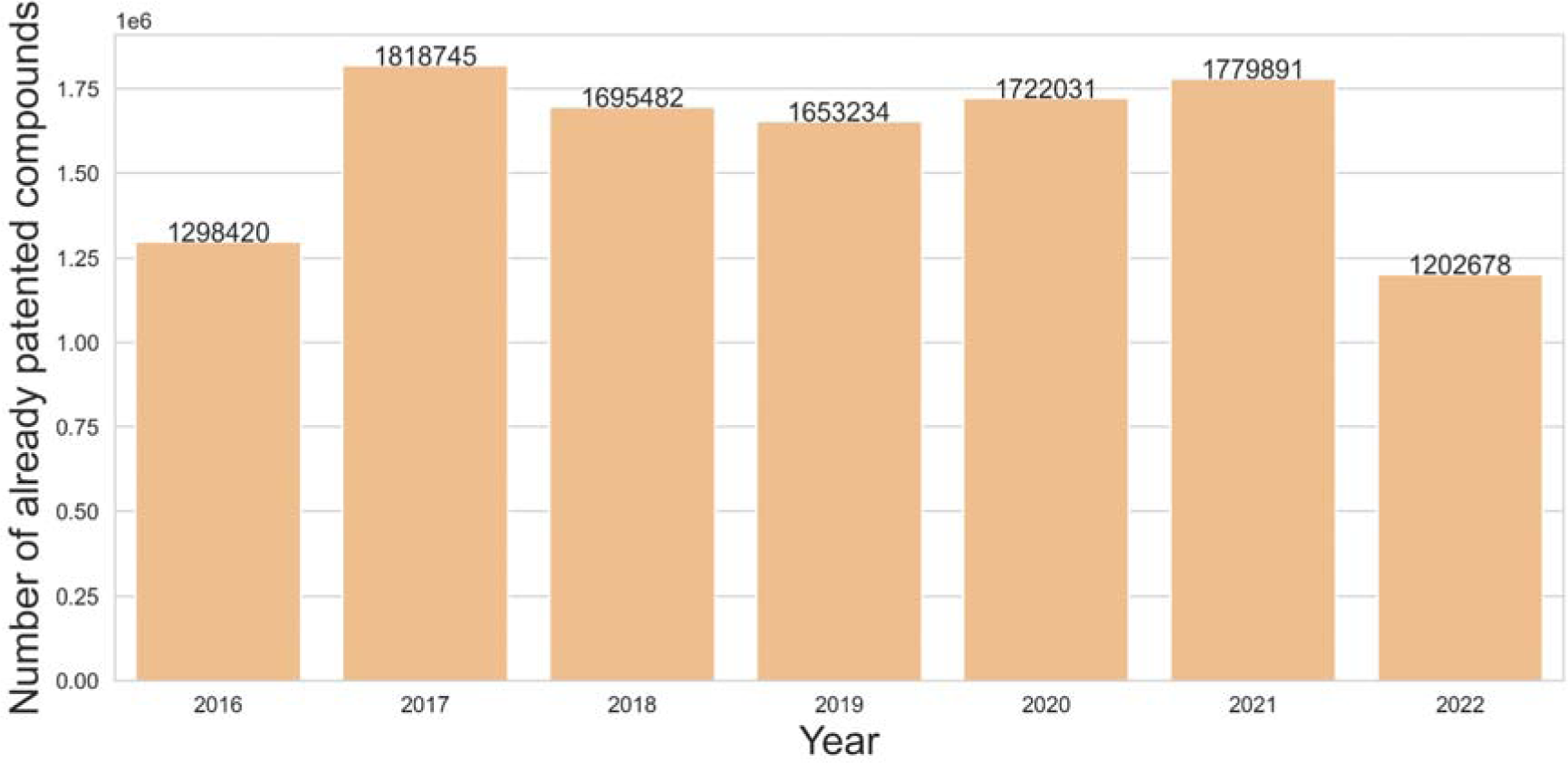
Repurposed patent compound distribution over the years. Distribution of patent compounds that have been found in patent documents from previous years. It is to be noted that the repurposing scenario shown here is only considered from 2015 onwards.

**Supplementary Figure SF2:**
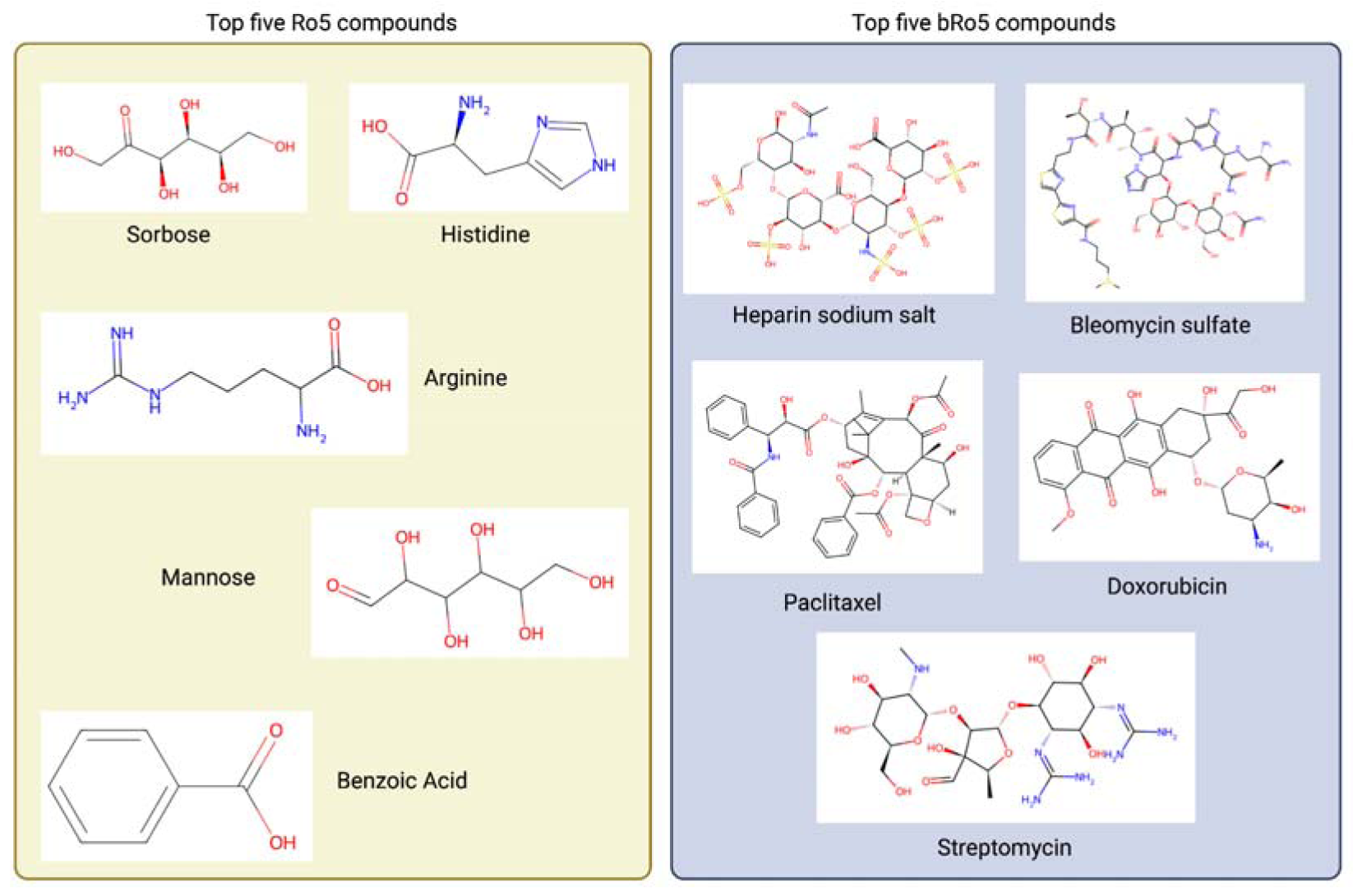
Top five prevalent patent compounds in the Ro5 and bRo5 categories.

**Supplementary Figure SF3:**
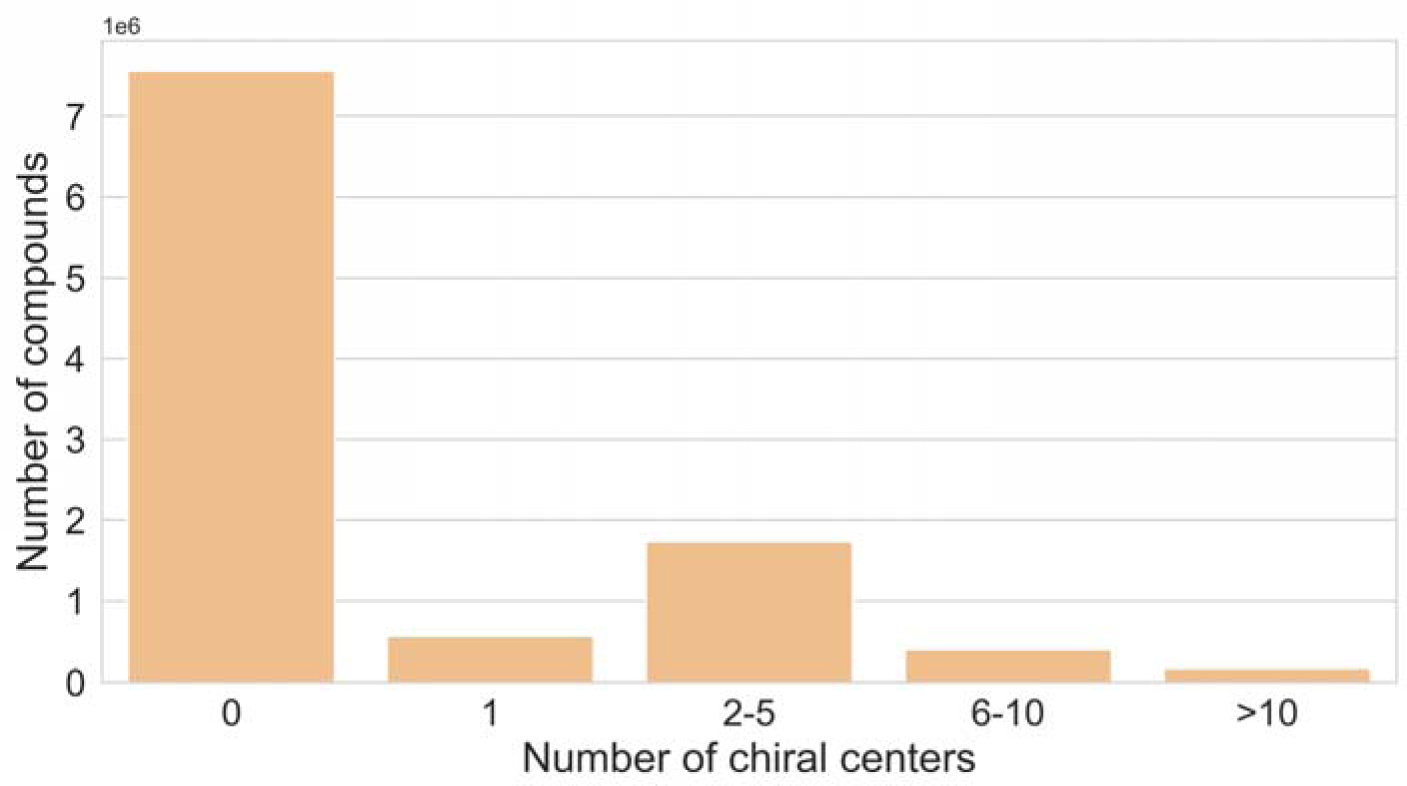
Distribution of the number of chiral centers across patent compounds.

**Supplementary Table ST1:** Summary of the number of compounds found with respect to patent kind and country of filing. The compounds were counted based on their unique InChIKey representation in SureChEMBL.

| Country | Patent kind code | Number of patent compounds |
| --- | --- | --- |
| United States (US) | Filed (A) | 6,111,699 |
|  | Granted (B) | 5,207,524 |
|  | Design Patent (S) | 2 |
|  | Reissue Patent (E) | 46,215 |
|  | Plant Patent (P) | 402 |
| Europe (EP) | Filed (A) | 2,648,617 |
|  | Granted (B) | 3,173,776 |
| Japan (JP) | Filed (A) | 254 |
|  | Granted (B) | 7 |

**Supplementary Table ST2:** Summary of the number of compounds found with patent documents. The compounds were counted based on their unique InChIKey representation in SureChEMBL.

| Number of patent documents | Number of compounds |
| --- | --- |
| < 5 | 10,148,500 |
| 10 - 50 | 445,227 |
| 50 - 100 | 37,247 |
| 100 - 500 | 37,157 |
| 500 - 1000 | 7,000 |
| > 1000 | 11,613 |

**Supplementary Table ST3:** Averaged yearly physicochemical property descriptions for compounds found in patent documents. For each year, we report the topological polar surface area, the number of potential stereoisomers and the number of heteroatoms.

| Year | TPSA | # Stereoisomers | # Heteroatoms |
| --- | --- | --- | --- |
| 2105 | 78.90 | 4.40 | 7.26 |
| 2016 | 82.64 | 5.05 | 7.66 |
| 2017 | 81.40 | 7.01 | 7.49 |
| 2018 | 80.43 | 6.57 | 7.56 |
| 2019 | 82.21 | 6.52 | 7.71 |
| 2020 | 82.94 | 6.47 | 7.80 |
| 2021 | 82.77 | 5.78 | 7.78 |
| 2022 | 81.79 | 6.37 | 7.83 |

**Supplementary Table ST4:**
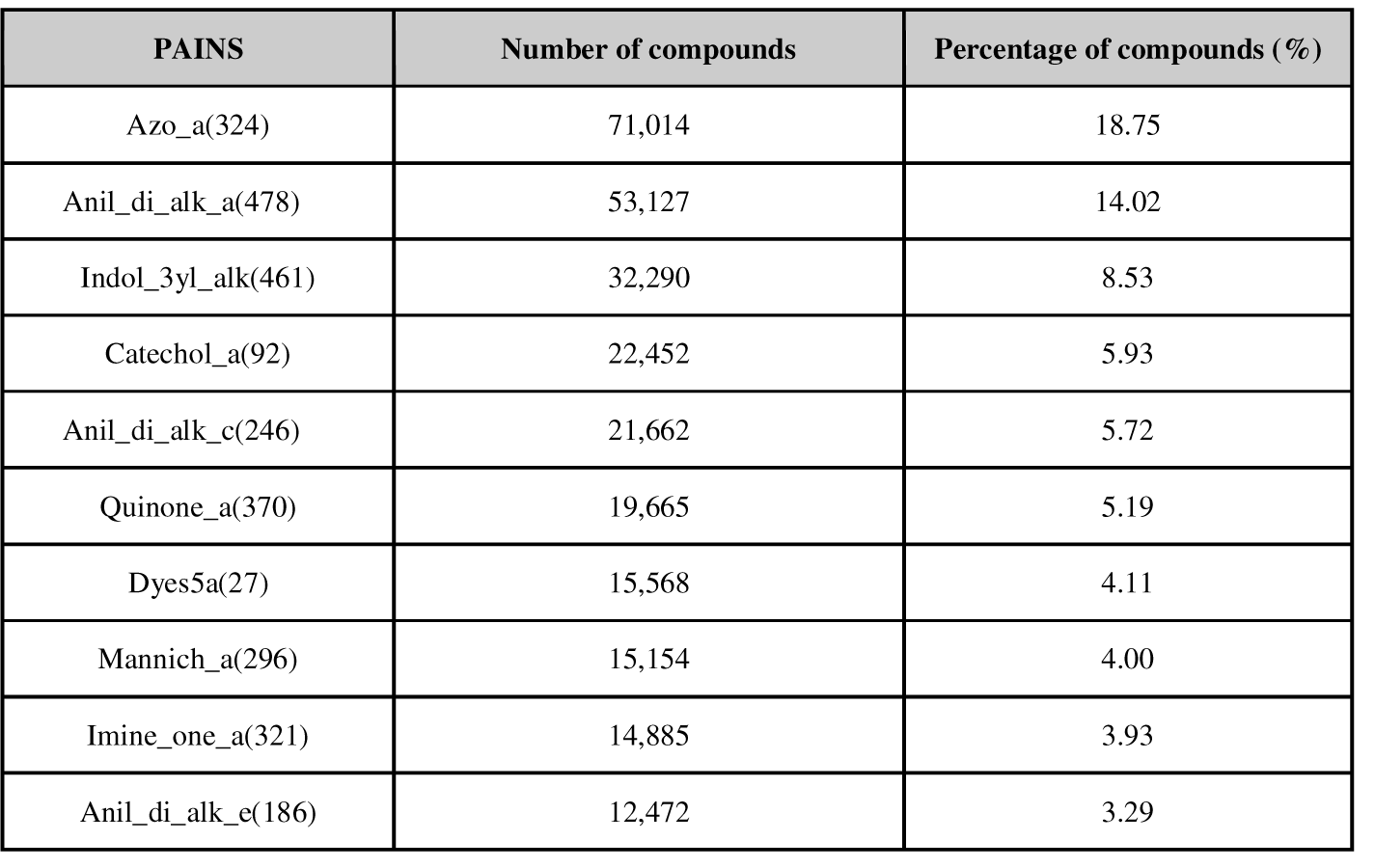
Summary of the top PAINS alerts found in patent compounds.

**Supplementary Table ST5:** Summary of the number of unique scaffolds found in the individual patent document sections.

**Supplementary Table ST4: Summary of the top PAINS alerts found in patent compounds.**
| Patent Field | Number of scaffolds | Percentage of scaffold (%) |
| --- | --- | --- |
| Abstract | 8,120 | 0.19 |
| Claims | 315,413 | 7.73 |
| Description | 811,216 | 19.89 |
| Image (after 2007) | 2,261,009 | 55.44 |
| MOL file (after 2007) | 677,947 | 16.62 |
| Title | 4,494 | 0.11 |

